# Greater resistance to footshock punishment in female C57BL/6J mice responding for ethanol

**DOI:** 10.1101/2022.08.05.502979

**Authors:** Elizabeth A. Sneddon, Kaila A. Fennell, Sachi Bhati, Joshua E. Setters, Kristen M. Schuh, Jenelle N. DeMedio, Brandon J. Arnold, Sean C. Monroe, Jennifer J. Quinn, Anna K. Radke

**Author notes:** Correspondence to:* Anna K. Radke, Ph.D., 90 N Patterson Ave, Oxford, OH, USA 45056.

## Abstract

**Background:** One characteristic of alcohol use disorder (AUD) is compulsive drinking, or drinking despite negative consequences. When quinine is used to model such aversion-resistant drinking, female rodents typically are more resistant to punishment than males. Using an operant response task where C57BL/6J responded for ethanol (EtOH) mixed with quinine, we previously demonstrated that female mice tolerate higher concentrations of quinine in EtOH than males. Here, we aimed to determine if this female vulnerability to aversion-resistant drinking behavior is similarly observed when footshock punishment is used.

**Methods:** Male and female C57BL/6J mice were trained to respond for 10% EtOH in an operant task on a fixed ratio 3 schedule. After consistent responding, mice were tested in a punishment session using either a 0.25 mA or 0.35 mA footshock. To assess footshock sensitivity, a subset of mice underwent a flinch, jump, vocalize test in which behavioral responses to increasing amplitudes of footshock (0.05 – 0.95 mA) were assessed. In a separate cohort of mice, males and females were trained to respond for 2.5% sucrose and responses were punished using a 0.25 mA footshock.

**Results:** Males and females continued to respond for 10% EtOH when paired with a 0.25 mA footshock. Females alone continued to respond for EtOH when a 0.35 mA footshock was delivered. Both males and females reduced responding for 2.5% sucrose when punished with a 0.25 mA footshock. Finally, footshock sensitivity in the flinch, jump, vocalize test did not differ by sex.

**Conclusions:** Females continue to respond for 10% EtOH despite a 0.35 mA footshock and this behavior is not due to differences in footshock sensitivity between males and females. These results suggest that female C57BL/6J mice are generally more resistant to punishment in an operant self-administration paradigm. These results add to the literature characterizing aversion-resistant alcohol drinking behaviors in females.

## Introduction

Alcohol use disorder (AUD) is characterized by behaviors such as compulsive alcohol drinking, or drinking that persists despite negative consequences (American Psychiatric Association, 2013). In patient populations, negative consequences such as problems with one’s health, finances, or relationships can have long-lasting impacts on wellbeing. Preclinical studies in rodents have sought to model this critical component of AUD by pairing ethanol (EtOH) delivery or a response for EtOH with an aversive stimulus (Hopf and Lesscher, 2014; Radke et al., 2021a). In one of the most commonly used models of aversion-resistant drinking, quinine, a bitter tastant, is added to the EtOH solution. We previously investigated whether male and female mice differ in their resistance to quinine punishment and found that females are more likely to respond for EtOH mixed with quinine than males (Sneddon et al., 2020). Interestingly, increased resistance to aversion in EtOH-drinking mice is not observed under all conditions (Radke et al., 2021a). For example, in a limited access, drinking in the dark task, male and female mice are equally sensitive to quinine punishment (Sneddon et al., 2019; Bauer et al., 2021). These disparate results point to the importance of characterizing how vulnerability for alcohol drinking behaviors varies across experimental parameters.

One critical experimental parameter in studies of aversion-resistant drinking is the aversive stimulus used to deter consumption or responding for EtOH. Some studies of aversion-resistant drinking have used a footshock punishment, typically delivered through the chamber floor when the animal makes a lever press or nose poke response in an operant chamber (Hopf and Lesscher, 2014). While both quinine and footshock serve as robust deterrents of EtOH responding, there are some important differences between the two. First, quinine is typically thought of as aversive due to its bitter gustatory properties while footshock aversion may additionally induce defensive responses. Second, quinine is delivered in the EtOH solution and therefore may serve to punish the licking response. Footshock is instead used to punish responses made in order to gain access to the EtOH solution. Thus, even when all other variables are held constant, there are differences between quinine and footshock punishment that could influence experimental outcomes.

Multiple studies have investigated the neural mechanisms driving aversion-resistant responding for EtOH using operant paradigms. Regions such as the medial orbitofrontal cortex (Radke et al., 2017), the medial prefrontal cortex (Halladay et al., 2020), the insula, and the nucleus accumbens (Seif et al., 2013; Chen and Lasek, 2020) have all been associated with this behavior in male rodents. While some studies using male rodents have demonstrated that quinine- and footshock-resistant EtOH drinking involve similar neural mechanisms (Seif et al., 2013; Siciliano et al., 2019), we do not know if this finding extends to females. Further, by using only male subjects, prior investigations leave a gap in our understanding of whether these paradigms are suitable to assess punished responding for EtOH in both sexes. To our knowledge, there have been no studies examining footshock-resistant responding for EtOH in female mice. As we have previously observed increased aversion-resistance in female mice responding for EtOH mixed with quinine, the experiments presented here were designed to assess whether this vulnerability is also observed using a footshock punishment.

To accomplish this goal, we adapted our previously validated operant response task to assess if footshock-punished responding for EtOH differs by sex. Based on previous findings (Sneddon et al., 2020), we hypothesized that females would show greater punishment-resistant responding despite a footshock while responding for 10% EtOH. In the first experiment, males and females continued to respond for 10% EtOH despite a 0.25 mA footshock. When a new cohort of mice was tested with a 0.35 mA footshock, females continued to respond for the 10% EtOH solution but responding in males was suppressed compared to baseline. In addition, we assessed whether males and females differed in footshock sensitivity but found no differences. In a second experiment, we assessed whether footshock would suppress responding for a non-drug reward. We found that a 0.25 mA footshock is sufficient to suppress footshock responses in male and female mice responding for a 2.5% sucrose solution. Collectively, these studies confirm that female vulnerability to aversion-resistant responding for EtOH in an operant paradigm persists despite punishment with a footshock.

## Methods

### Subjects

Fifty-one adult male (n = 25) and female (n = 26) C57BL/6J mice were bred from breeding pairs purchased from Jackson Laboratory (Bar Harbor, ME) and housed in the Laboratory of Animal Resources at Miami University. Mice had access to Rodent Diet 5001 chow (Cincinnati Lap Supply, Cincinnati, OH) and reverse-osmosis (RO) drinking water *ad libitum*, unless noted otherwise. Mice were kept in a temperature-controlled room on a 12:12 hour light:dark cycle (lights on at 7:00 AM). Prior to and throughout operant training, mice were housed in groups of 2 – 3 per cage. Each cage was a standard shoe box udel polysulfone rectangular mouse cage (18.4 × 29.2 × 12.7 cm) with 5.08 × 5.08 cm nestlets (Cincinnati Lab Supply, Cincinnati, OH), and Bed-O-Cob 0.64 cm bedding (Cincinnati Lab Supply, Cincinnati, OH). All subjects were cared for in agreement with the guidelines set by the National Institutes of Health and all procedures were approved by the Institutional Animal Care Use Committee (IACUC) at Miami University.

### Drinking solutions

EtOH (10%) solution was prepared volume/volume in RO water. Sucrose (10%) solution was prepared weight/volume in RO water. For the 10% sucrose + 10% EtOH and 5% sucrose + 10% EtOH solutions, sucrose was prepared weight/volume before being added to 10% EtOH. Sucrose (2.5%) solution was prepared weight/volume in RO water. All EtOH solutions were made fresh prior to each testing session. Sucrose solutions without EtOH were prepared weekly and stored at 20°C.

### Operant apparatus for EtOH responding

A standard 15.24 × 13.34 × 12.7 cm mouse conditioning chamber was used for all operant training (Med Associates, Fairfax, VT, ENV-307A) (as described in Sneddon et al., 2020). The chamber had a grid floor with nineteen rods and 0.79 cm of space between each rod (ENV-307A-GFW). The mouse conditioning chamber was housed within a sound and light attenuating box (ENV-022 V). A house light was on one wall of the chamber and parallel to the wall of the chamber there was a reward receptacle (303RMA-3) and two nose poke ports (ENV-313 M). The left port was designated the active port and the right port designated as the inactive port during all training. For training involving grain pellets (14 mg), a pedestal mount pellet dispenser (ENV-203-20) was attached to the reward receptacle. For sessions where liquid solutions were dispensed (50 µL over 1.5 s), a single speed syringe pump (PHM-100) with a 20-mL syringe was used. At the beginning of each session, the light above the reward receptacle (ENV-303RL) was illuminated and stayed on throughout the duration of the session. The main house light was off throughout each session. A 2 s, 65-db tone sounded when a reward was delivered (as described in Radke et al., 2017b). To deliver a footshock to the mice, an aversive stimulator/scrambler (ENV-414S) was used. A shock was delivered on the second nose poke response for 0.5 sec. After each test session, the chamber was cleaned with 70% EtOH and syringes that contained the solutions were cleaned with RO water.

### Response training for EtOH

Two weeks prior to testing, mice were food restricted to 85% of their free feeding weight to increase engagement with the task (Rowland, 2007). Throughout the operant phases of the experiment, mice were kept on food restriction and were fed once daily following each testing session. The amount of food (1.8 – 4.0 g/mouse/day; Rodent Diet 5001 chow) was adjusted daily to ensure that all mice maintained 85% of their free feeding weight. Behavioral testing occurred 3 – 6 h into the light cycle (*i*.*e*., 10 AM – 1 PM) on Mondays - Fridays. Mice were fed within this timeframe on the weekends to maintain their food restriction.

First, mice were trained to nose poke on a Fixed-Ratio (FR) 1 schedule on the active nose port for a grain pellet in 30 min for 3 sessions. During the next session mice underwent a “sucrose-fading” procedure where they had to respond first for 10% sucrose on an FR1 schedule for 3 sessions before transitioning to a FR3 schedule for 3 – 5 sessions pending response stabilization (coefficient of variation < 20% across 3 consecutive sessions). When responding stabilized or once mice met the maximum of 5 sessions, EtOH was added to the sucrose solution on the next session. The sucrose was faded out in the following concentrations: 10% sucrose + 10% EtOH, 5% sucrose + 10% EtOH, and 10% EtOH (Sneddon et al., 2020, Halladay et al., 2017, Radke et al., 2017). Mice responded for each solution for 3 – 5 sessions and then remained on 10% EtOH until their responses stabilized (as described above). Drinking cups were checked after each session to verify consumption.

### Punished EtOH responding

To assess if mice would continue to respond despite the punishment of a footshock, two separate cohorts of mice were used. The first cohort (n = 16, male = 8, female = 8) underwent the response training described above. On the session following stabilization of responding, footshock punishment commenced using a 0.25 mA footshock. The second cohort (n = 17, male = 9, female = 8) underwent the same experimental procedure except their responses were punished with a 0.35 mA footshock (**Fig. 1A**). During punishment sessions, mice responded on an FR3 schedule and footshock was always delivered on the second nose poke response (after Radke et al., 2017b).

**Figure 1.**
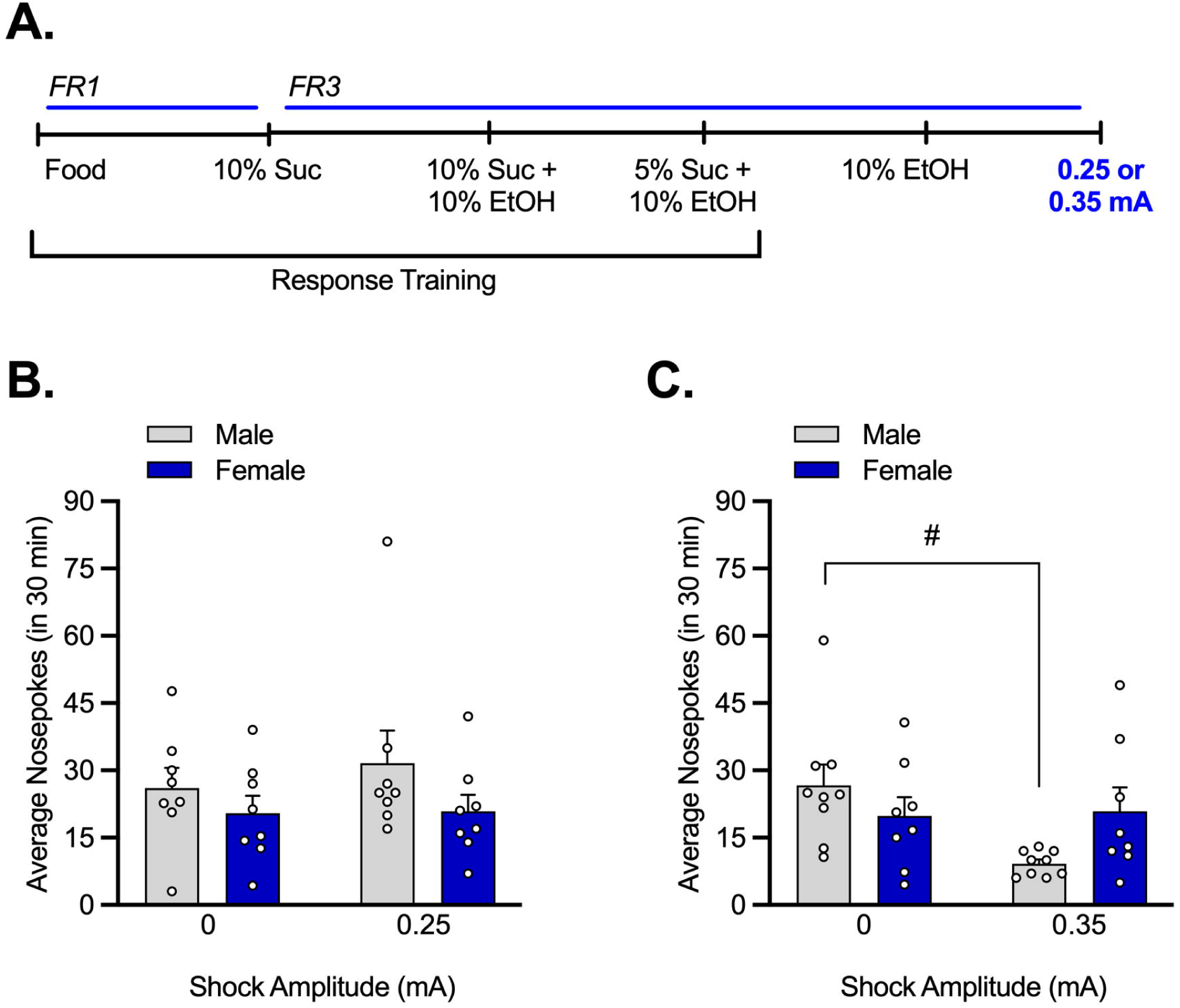
Females show greater resistance to a footshock when responding for EtOH. (A) Response training consisted of the “sucrose fading” procedure (grain pellets → 10% sucrose → 10% sucrose + 10% ethanol (EtOH) → 5% sucrose + 10% EtOH → 10% EtOH) followed by a session where a 0.25 mA or 0.35 mA footshock was delivered during one 10% EtOH session. (B) Males and females continue to respond for EtOH despite a 0.25 mA footshock but (C) only females continue to respond despite a 0.35 mA footshock. # *p* < 0.05 vs. 0 mA (Holm-Sidak’s).

### Flinch, jump, vocalize test

At least two weeks following response training, a subset of mice who were punished with the 0.25 mA shock (n = 14, male = 7, female = 7) underwent a footshock sensitivity test using the flinch, jump, vocalize paradigm (modified after Kim et al., 1991). Flinch was operationalized as an observable reaction to the shock (lifting or shaking their paws or directing their attention toward the grid floor). Jump was defined as any jumping, prancing, or running following the footshock. Vocalize was operationalized as an audible squeak when shocked (Kim et al., 1991).

A 32.4 × 25.4 × 21.6 cm^3^ operant chamber housed in a sound attenuating chamber was used for the footshock sensitivity test. The chamber consisted of a grid floor and house light on one wall. The chamber was cleaned with odorless 5% sodium hydroxide following each test session. Grid floors were connected to a shock generator and scrambler (as previously described in Quinn et al., 2014; Radke et al., 2020; Sneddon et al., 2021). First, mice were placed in the chamber and were allowed to explore for 3 min. At 3 min, a 0.05 mA shock was delivered. Mice were then shocked every minute in increasing amplitudes (from 0.05 mA – 0.95 mA). Following the last shock, mice remained in the chamber for 1 min, for a total of 13 min 10 s. The researcher manually increased the shock amplitude between footshocks. Each mouse was tested and monitored individually and the experimenter logged the first instance of the flinch, jump, and vocalize responses.

### Punished sucrose responding

To determine if punished responding was specific to the EtOH reward, naïve mice (n = 17, male = 8, female = 9) were first trained to respond for a grain pellet on an FR1 schedule in 30 min across 3 sessions (all training occurred in 30-min sessions). Mice were then trained to respond for 2.5% sucrose on an FR1 schedule for at least 3 sessions or until their responding stabilized (as described above) before transitioning to an FR3 schedule. Once responding stabilized, responding for sucrose was punished with a 0.25 mA footshock on one session (**Fig. 2A**).

**Figure 2.**
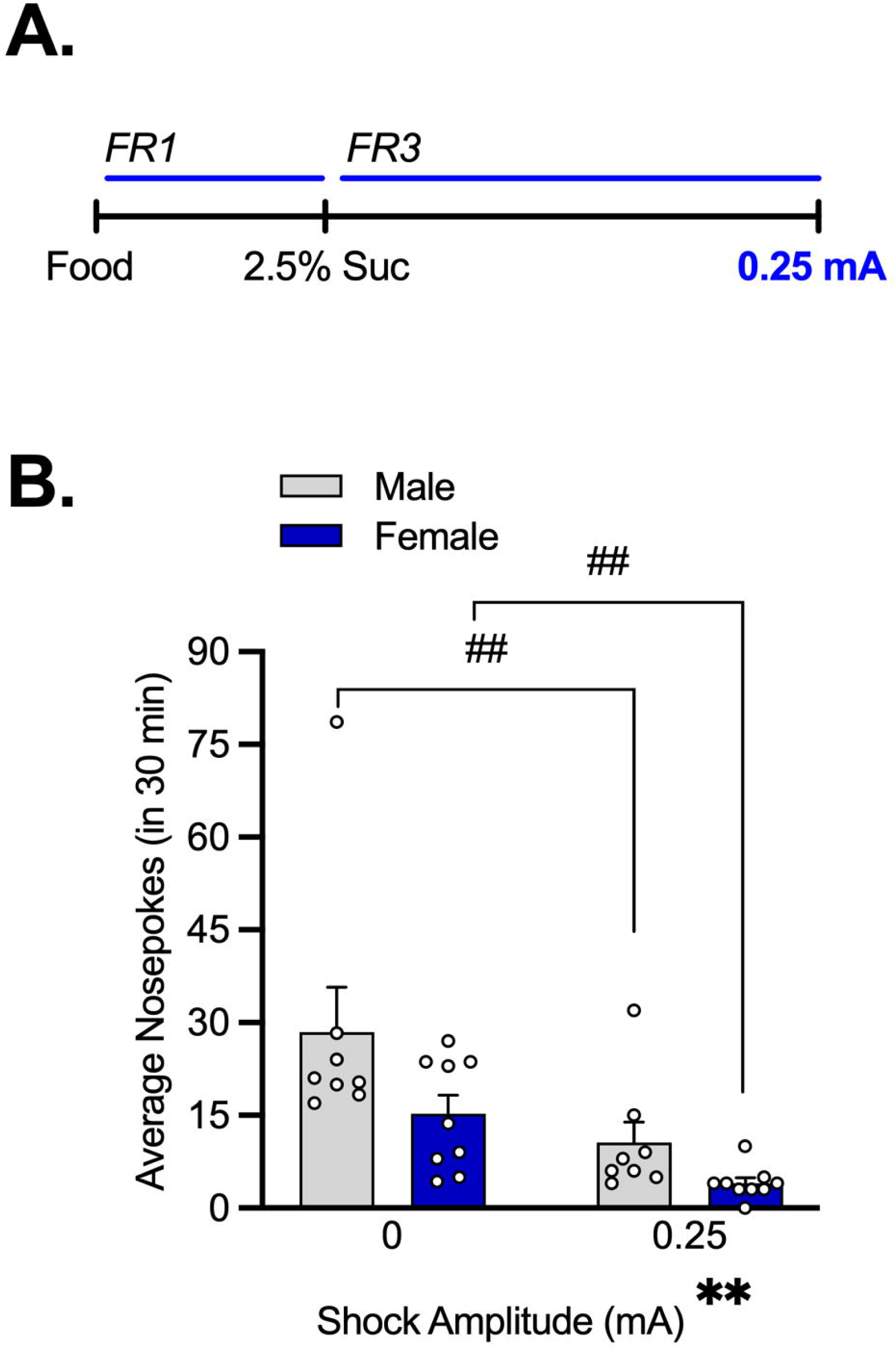
A 0.25 mA footshock suppresses sucrose seeking equally in males and females. (A) Mice were trained to respond for 2.5% sucrose on an FR3 schedule and then responses were paired with a 0.25 mA footshock during one session. (B) Responses were reduced in both sexes when sucrose delivery was paired with a footshock. ## *p<* 0.01 vs. 0 mA (Holm-Sidak’s) and ** *p<* 0.01 main effect of shock amplitude (2-Way ANOVA).

### Data Analysis

Nose poke responses were calculated as the number of nose pokes made per session. Responses were averaged across the last three sessions of response training to calculate baseline responding. Flinch, jump, vocalize values were calculated as averages of shock amplitude across subjects.

Responses for EtOH or sucrose were analyzed using a Two-Way Repeated-Measures (RM) Analysis of Variance (ANOVA) with sex as the between subject-factor and solution or shock amplitude as the within-subjects factor. *Post hoc* Holm Sidak’s tests were planned to make comparisons between baseline vs. punished responding for each sex. Footshock sensitivity results were analyzed using a Two-Way RM ANOVA with sex as the between subject-factor and behavioral response (flinch, jump, vocalize) as the within-subjects factor. In cases where the assumption of sphericity was violated (ε < 0.75), the Greenhouse-Geisser correction was used. All data were expressed as mean ± standard error of the mean and was analyzed using GraphPad Prism v. 9.0 (La Jolla, CA).

## Results

### Females show greater resistance to a footshock when responding for 10% EtOH than males

#### Response training

Males and females completed a similar number of sessions throughout response training for 10% sucrose + 10% EtOH, 5% sucrose + 10% EtOH, and 10% EtOH (**Table 1)**. For mice in the 0.25 mA cohort, A Two-Way RM ANOVA found a main effect of solution (F_(1.067, 14.932)_ = 6.981, p = 0.017) but no main effect of sex (F_(1, 14)_ = 0.000, p > 0.999) and no interaction (F_(2, 28)_ = 0.292, p = 0.749) (**Table 1**). When assessing the amount of EtOH delivered, a Two-Way RM ANOVA showed a main effect of solution (F_(1.624, 22.733)_ = 15.581, p = 0.0001) but no main effect of sex (F_(1, 14)_ = 0.288, p = 6.00) and no interaction (F_(2, 28)_ = 0.741, p = 0.486) (data not shown). For mice in the 0.25 mA cohort, the amount of 10% EtOH delivered at baseline was 1.394 ± 0.223 g/kg for males and 1.047 ± 0.1 g/kg for females [data expressed as mean ± standard error of the mean].

**Table 1.**
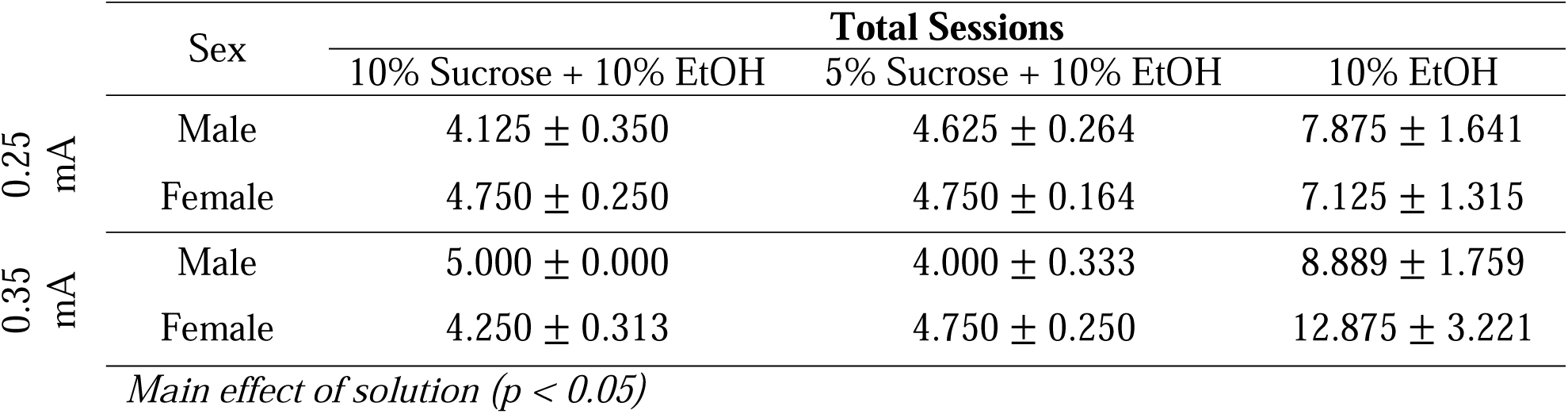
Total sessions during response training. Male and female mice completed the same number of total sessions during each phase of response training. Total sessions varied by solution. Data expressed as averages ± standard error of the mean (SEM). *\*p < 0.05* (2-Way ANOVA).

Similarly, mice in the 0.35 mA cohort completed a similar number of sessions throughout response training, a RM Two-Way ANOVA revealed a main effect of solution (F_(1.023, 15.346)_ = 13.145, p = 0.002) but no main effect of sex (F_(1, 15)_ = 1.140, p = 0.303) and no interaction (F_(2, 30)_ = 1.417, p = 0.258) (**Table 1**). When assessing the amount of EtOH delivered, a Two-Way RM ANOVA identified a main effect of solution (F_(1.870, 28.054)_ = 25.678, p < 0.0001) but no main effect of sex (F_(1, 15)_ = 0.061, p = 0.808) and no interaction (F_(2, 30)_ = 1.725, p = 0.195) (data not shown). For mice in the 0.35 mA cohort, at baseline, males received 1.520 ± 0.249 g/kg and females received 1.291 ± 0.312 g/kg of 10% EtOH.

#### Responses for EtOH

Males and females continued to respond for a 10% EtOH reward when punished with a 0.25 mA footshock. A Two-Way RM ANOVA identified no main effects of sex (F_(1, 14)_ = 2.094, p = 0.170) or shock amplitude (F_(1, 14)_ = 0.472, p = 0.504) and no interaction (F_(1, 14)_ = 0.338, p = 0.370) (**Fig. 1B**). When assessing the amount of EtOH delivered, a Two-Way RM ANOVA found no main effects of sex (F_(1, 14)_ = 0.093, p = 0.764) or shock amplitude (F_(1, 14)_ = 0.171, p = 0.686) and no interaction (F_(1, 14)_ = 0.796, p = 0.387) (data not shown). During the shock session, males received 1.655 ± 0.430 g/kg and females received 1.247 ± 0.279 g/kg of 10% EtOH.

Females made more responses for 10% EtOH despite a 0.35 mA footshock than males. A Two-Way RM ANOVA revealed no main effects of sex (F_(1, 15)_ = 0.402, p = 0.535) or shock amplitude (F_(1, 15)_ = 3.553, p = 0.079). The interaction between sex and shock amplitude approached the threshold for significance (F_(1, 15)_ = 4.534, p = 0.0502). A *post hoc* Holm Sidak’s test found that males made significantly fewer responses for 10% EtOH when responses were paired with a 0.35 mA shock (p = 0.021) but responding for EtOH was maintained in females (**Fig. 1C**). When assessing the amount of EtOH delivered, a Two-Way RM ANOVA identified a main effect of shock amplitude (F_1,15)_ = 4.659, p = 0.048) and no main effect of sex (F_(1,15)_ = 1.169, p = 0.297) and no interaction (F_(1,15)_ = 3.176, p = 0.095). A *post hoc* Holm Sidak’s test showed that males earned less 10% EtOH when responses were paired with a 0.35 mA shock (p = 0.023) but the amount of EtOH delivered was maintained in females (data not shown). During the shock session, males received 0.444 ± 0.606 g/kg and females received 1.189 ± 0.341 g/kg of 10% EtOH.

### Footshock suppresses sucrose responding equally in males and females

#### Response training

Males and females completed a similar number of sessions throughout response training. An unpaired t-test found no difference in the number of 2.5% sucrose training sessions prior to shock exposure (t_(15)_ = 1.775, p = 0.096) for males (= 7.222 ± 0.813) or females (=5.625 ± 0.263) (data expressed as mean ± standard error of the mean).

#### Responses for sucrose

Footshock (0.25 mA) suppressed responses for 2.5% sucrose in both sexes. A Two-Way RM ANOVA revealed a main effect of shock amplitude (F_(1, 15)_ = 28.471, p < 0.0001) but no main effect of sex (F_(1, 15)_ = 3.735, p = 0.072) and no interaction (F_(1, 15)_ = 1.453, p = 0.247). A *post hoc* Holm Sidak’s test found that both males and females made fewer responses for sucrose when paired with a 0.25 mA footshock (p < 0.01) (**Fig. 2B**).

### Males and females do not differ in footshock sensitivity

At least two weeks following punished EtOH responding, a subset of mice from the 0.25 mA cohort (n = 14, male = 7, female = 7) underwent a footshock sensitivity test to assess sex differences. Flinch, jump, and vocalization responses were induced at different shock amplitudes, but the responses did not differ by sex. A Two-Way RM ANOVA identified a significant main effect of response type (F_(2,24)_ = 72.488, p < 0.0001) but no main effect of sex (F_(1,12)_ = 0.565, p = 0.4669) and no interaction (F_(2,24)_ = 0.488, p = 0.6196) (**Table 2**).

**Table 2.** Footshock sensitivity does not differ by sex. Males and females show observable flinch, jump, or vocalize behaviors at similar shock amplitudes. Data expressed as averages ± standard error of the mean (SEM).

| Response | Male | Female |
| --- | --- | --- |
| Flinch | $0.107 \pm 0.020$ | $0.107 \pm 0.020$ |
| Jump | $0.221 \pm 0.018$ | $0.207 \pm 0.020$ |
| Vocalize | $0.393 \pm 0.037$ | $0.350 \pm 0.031$ |

## Discussion

The major finding of this work is that in an operant self-administration paradigm, male and female C57BL/6J mice show similar footshock-resistant responding for 10% EtOH when the amplitude is set to 0.25 mA, but females alone continue to exhibit footshock-resistant responding when the amplitude is increased to 0.35 mA. In addition, footshock-resistant responding is specific to an EtOH reinforcer as both sexes reduce their responding for 2.5% sucrose that is paired with a 0.25 mA footshock. Lastly, male and female mice do not differ in their sensitivity to footshock, as evidenced by similar response thresholds during a flinch, jump, vocalize test. These results suggest that C57BL/6J female mice are more motivated to respond for an EtOH reward despite the negative consequence of a footshock.

In two separate cohorts of mice, we tested punishment-resistant responding for 10% EtOH. As in our previous study (see Sneddon et al., 2020), sex differences in responding for 10% EtOH were not observed during response training or baseline sessions in either cohort. The use of a 10% EtOH solution here allows direct comparison of the current results with our previous investigations of quinine-resistant responding for EtOH. Similar response rates between males and females for this concentration of EtOH also ensure that differences in footshock-resistant responding are not due to differences in the levels of EtOH exposure during training, as this factor is known to influence the development of punishment-resistance (Radke et al., 2017; Houck et al., 2019; Sneddon et al., 2020).

The current results demonstrate that both males and females continue to respond for 10% EtOH when a 0.25 mA footshock is paired with the nose-poke response. This level of footshock was sufficient, however, to suppress responding for 2.5% sucrose in both sexes. Together, these findings demonstrate that both males and females demonstrate footshock-resistant responding for EtOH in the operant response task. This result is not surprising given that we have previously found evidence for quinine-resistant responding for 10% EtOH in both sexes using this same behavioral paradigm (Sneddon et al., 2020). We have also demonstrated that male C57BL/6J mice are resistant to footshock punishment when responding for EtOH vs. a food pellet once before, although the levels of footshock required to suppress responding for reward were higher in that study (Radke et al., 2017b). Thus, punishment resistant responding for EtOH is a reliable finding in this strain of mice.

Although both sexes demonstrated footshock-resistance at the lower shock amplitude, when the amplitude was increased to 0.35 mA in a new cohort of mice females continued to respond for 10% EtOH while responding in males was suppressed. It is important to note that while our conclusions regarding the effects of the 0.35 mA footshock are supported by planned comparisons between the baseline and footshock sessions, the interaction of sex X shock amplitude was just above the threshold for significance (p = 0.0502), raising the possibility that this portion of the study was slightly underpowered. The robustness of our findings are increased, however, when considered alongside our previous work on quinine-resistant responding for EtOH (Sneddon et al., 2020). Using the identical operant paradigm described here, we observed suppression of responding in males at quinine concentrations of 250 and 500 µM but no change in female responding for 10% EtOH at any quinine concentration (Sneddon et al., 2020). Together, these results suggest that female C57BL/6J mice have a higher tolerance for punishment when responding for EtOH (but not sucrose) in this operant paradigm.

In addition to verifying that the effects seen here are specific to EtOH reward by testing a separate cohort of mice responding for sucrose, we also controlled for the possibility that punishment-resistance could be driven by sex differences in footshock sensitivity. Sensitivity to footshock was tested in male and female C57BL/6J mice using a flinch, jump, vocalize test (Kim, 1991). Behavioral responses to footshock were similar in males and females across the range of shock amplitudes, in agreement with prior reports (Podhorna et al., 2002). Again, this result is reminiscent of our work with quinine punishment where males and females avoid similar concentrations of quinine in water (Sneddon et al., 2019; Sneddon et al., 2020). Controls such as these are important as they support the conclusion that sex differences in punishment-resistance are driven by the motivational properties of EtOH and not due to differences in punishment sensitivity.

One limitation of this study is that the design of the operant task does not permit precise measurement of consumption. Rather than providing access to a sipper, completing the response requirement resulted in delivery of 50 µL EtOH into a cup. To verify consumption, drinking cups were checked at the end of each session. It is also important to consider that the observed results may be specific to the experimental parameters employed here. For example, female rats have been shown to respond more than males for 10% EtOH when an FR1 schedule of reinforcement is used (Flores-Bonilla et al., 2021). Interestingly, in agreement with our own work, this sex difference was not observed with an FR3 schedule of reinforcement. The concentration of EtOH may also be important, as we have shown that females respond more than males when higher concentrations of EtOH are delivered (Sneddon et al., 2020).

There has been a recent surge of interest in characterizing aversion-resistant drinking in male and female rodents across different paradigms (Radke et al., 2021b). Although the current study is the first to characterize sex differences in footshock-resistant responding in an operant task, one study examining EtOH conditioned place preference (CPP) found that experience with a footshock on the EtOH-paired side reduced preference in male but not female mice (Xie et al., 2019). Our finding that females maintain responding for EtOH despite a footshock while males cease responding agrees with this result. In home cage paradigms, the observance of sex differences in aversion-resistant responding has been more varied. For example, in a continuous access paradigm, female mice tolerated higher concentrations of quinine compared to males (Fulenwider et al., 2019) whereas in a DID paradigm male and female mice exhibited similar aversion-resistant EtOH drinking (Bauer et al., 2021; Sneddon et al., 2019). Another study employing a continuous access paradigm in which mice had access to water, nicotine, or 20% EtOH found that both sexes reduced their consumption of quinine-adulterated EtOH to similar degrees (DeBaker et al., 2019). In addition, we have shown that female rats with a history of early life stress are more vulnerable to quinine-resistant drinking compared to males and compared to females who did not experience early life stress (Radke et al., 2020). Taken together, while there appears to be a female vulnerability to aversion-resistant EtOH drinking, this sex difference emerges under some but not all conditions.

In summary, these findings show that female vulnerability to footshock-resistant responding can be studied in mice using an operant paradigm. Female mice continued to respond more for EtOH despite a 0.35 mA footshock while both sexes continued to respond for EtOH despite a 0.25 mA footshock. These results are specific to EtOH as both sexes reduced their responding for 2.5% sucrose when paired with a 0.25 mA footshock. In addition, both sexes exhibit similar behavioral responses to varying footshock amplitudes. This operant paradigm, with quinine or footshock punishment, can be useful for investigating mechanisms that may drive female vulnerability to EtOH drinking behaviors.

## Acknowledgments

The authors would like to thank Natalie Shand and Julia Hoffman for assistance with behavioral experiments.

